# A single binge ethanol exposure is apoptotic within hours across neurodevelopment and partially regulated by the *Myt1l* gene

**DOI:** 10.1101/2025.06.27.662020

**Authors:** Nicole Fuhler, Cory Palmer, Gianna Tunzi, Lilly Tian, Maya Fotedar, Rebecca Chase, Ramachandran Prakasam, Susan E. Maloney, C. Eric Neblock, Jiayang Chen, Kristen L. Kroll, Joseph D. Dougherty, Kevin K. Noguchi

## Abstract

Ethanol rapidly produces widespread neuronal apoptosis during early development, but this susceptibility declines as the brain matures. In previous research, we found *Myt1l* (a proneuronal transcription factor) mutations can cause precocious differentiation, neuronal immaturity, and transcriptomic alterations, including many in apoptotic regulators. Therefore, we used a recently developed *Myt1l* haploinsufficient mouse model to examine this gene’s effects on ethanol-induced apoptosis across different developmental stages. We discovered that haploinsufficiency can moderately influence vulnerability to ethanol in a complex, age- and cell type-specific manner: apoptosis was reduced on P7, increased P21, but unaffected on P60. Remarkably, we also discovered the previously unrecognized ability of a single binge of ethanol to rapidly increase apoptosis within six hours in early adolescent and adult wild-type mice occurring in microglia and the newborn granule neurons in the hippocampus. This suggests apoptosis is an underappreciated contributor to ethanol’s neuropathology at older ages and, translated to human use, occurs far more frequently than previously recognized.

## Introduction

The neurotoxicity produced by ethanol is highly age dependent. For instance, fetal alcohol spectrum disorders (FASD) are a group of physical, behavioral, and cognitive disorders caused by prenatal ethanol exposure^1,2^. Using FASD animal models, researchers have discovered a window of vulnerability called the ‘synaptogenesis period’ where ethanol exposure causes widespread neuronal apoptosis by inhibiting the brain through N-methyl-D-aspartate (NMDA) antagonism and GABA_A_ agonism^3,4^. In the mouse, the synaptogenesis period begins at the end of the third trimester, peaks around postnatal day 7 (P7), and ends around P14 (this extends from the end of the second trimester until several years after birth in the human)^4,5^. During normal development, the brain overproduces neurons but eliminates those that fail to make beneficial synaptic connections through apoptosis. Ethanol’s inhibition is thought to simulate poor synaptic connectivity and artificially induce apoptosis in healthy neurons^6^. After the synaptogenesis period, neurotoxicity studies primarily focus on more chronic ethanol exposures often lasting days or weeks, but its underlying causes are poorly understood. In humans, prolonged use can produce alcohol related brain damage (ARBD) — an umbrella term used to describe a set of related conditions caused by *chronic* ethanol use that includes secondary factors such as nutritional deficiencies, liver toxicity, and other compensatory mechanisms in the brain^7–9^. Over 20 years ago, chronic ethanol exposure in rodents was found to produce neurodegeneration in the dentate gyrus (DG) and olfactory related regions such as the entorhinal, piriform, and perirhinal cortices^10,11^. Unfortunately, these more prolonged regimens can complicate interpretation when trying to determine how repeated use becomes chronically neurotoxic and to what extent neurotoxicity is attributable to ethanol itself versus secondary factors. Indeed, even today questions remain about how this neuropathology occurs with suggested mechanisms including (among others): compensatory stimulation producing excitotoxicity, oxidative stress due to ethanol metabolism, edema, and neurotoxicity resulting from microglial activation and inflammation^12–16^.

While brain maturity is important, genetic factors also play an important role in ethanol’s neurotoxicity. Only 4.3% of children heavily exposed to ethanol will develop the most severe form of FASD. Additionally, two twin studies have found FASD diagnoses were 100% concordant for monozygotic twins but far less (64% and 56%) for dizygotic twins^17–20^. While several genes are associated with FASD, many play a role in ethanol metabolism with the others remaining poorly understood^18,21,22^. Genetics may also play a role in ARBD after the synaptogenesis period with research suggesting neurotoxicity is influenced by genes that affect thiamine deficiency or the function of NMDA or GABA_A_ neurotransmission^23–25^. Since ethanol is more neurotoxic in the immature brain, genes that affect neuronal development or apoptosis may also influence vulnerability. MYT1L is a proneuronal transcription factor highly expressed in postmitotic neurons which is sustained at low levels throughout life^26^. Human mutation in the *MTY1L* gene leads to a MYT1L Neurodevelopmental Syndrome (MNS) characterized by a variety of challenges, including developmental, language, and motor delays, as well as increased prevalence of obesity, seizure, ADHD, and autism. In previous research, we generated mice with a patient-inspired stop-gain mutation resulting in complete protein loss^27^. While *Myt1l* homozygous mutants die at birth, *Myt1l* heterozygous mice (Hets), which match the human patient genotype, exhibit a decrease in transcripts and protein. Het mice also exhibit precocious differentiation of neural progenitor cells into neurons but, even in adulthood, exhibit signs of neuronal immaturity including reductions in mature neuronal markers and immature spine morphology^27–29^. Consistent with this, generation of iSPC cellular models of MNS have revealed dysregulated neuronal maturation trajectories, and for one of them, a precocious progenitor exit from cell cycle consistent with our mouse model^30,31^.

Here we describe a reanalysis of existing iPSC data which suggests MYT1L haploinsufficiency dysregulates apoptosis. These data, and MYT1L’s ability to affect brain maturation throughout the lifespan, prompted us to examine this gene’s acute effects on ethanol-induced apoptosis in neonates, early adolescents, and adults (P7, P21, and P60 respectively). We discovered that haploinsufficiency can moderately influence vulnerability in a complex age- and cell type-specific manner with apoptosis reduced in neurons at P7, increased in microglia at P21, and unaffected at P60. Remarkably, we also discovered the previously unknown ability of a single binge of ethanol to rapidly produce apoptosis within 6 hours in microglia and newborn dentate gyrus granule neurons during the early adolescent and adult periods. This indicates that apoptosis is an underappreciated aspect of ethanol’s neurotoxicity after the synaptogenesis period and may occur far more frequently than previously suspected during human use.

## RESULTS

### MYT1L haploinsufficiency transcriptionally alters apoptotic regulators in differentiating neurons

MYT1L haploinsufficiency has widespread consequences due to its role as a proneural transcription factor, especially in pure populations of differentiating neurons *in vitro*. We set out to investigate these consequences using previously published hPSC RNA sequencing (RNAseq) data comparing an engineered variant knockin model associated with MYT1L haploinsufficiency (VKI) to a wild type (WT) hPSC model^32^. We focused on robustly differentially expressed genes (DEGs): i.e., with an FDR-adjusted p-value less than 0.05 and a log2 fold change that was either greater than 0.5 or less than −0.5. This resulted in 472 genes classified as upregulated and 1336 downregulated genes in the VKI model (**Figure 1A**). We then investigated the biological processes that may be impacted by MYT1L haploinsufficiency. Interestingly, when looking at processes associated with downregulated genes, we find that many have to do with apoptosis. Notably, the downregulated genes were associated with both positive and negative regulation of apoptosis (**Figure 1B**). Thus, while it is clear there is some transcriptional dysregulation of apoptotic programs due to MYT1L loss, it was unclear if MYT1L loss might promote or suppress apoptosis. To further confirm this association, we performed a gene set enrichment analysis (GSEA) on the full set of ranked log2 fold changes, using the gene set defined by the molecular signatures database (MSigDB) as hallmark apoptosis genes. The results of this analysis suggest that hallmark apoptosis genes are downregulated in this VKI model of MYT1L haploinsufficiency, with a normalized enrichment score (NES) of −1.58 and an adjusted p-value of 0.0026 (**Figure 1C**). Further investigation into individual genes confirmed that a majority of these hallmark apoptosis genes were downregulated in the VKI model. However, there do exist some hallmark apoptotic genes that appear to be upregulated in the VKI model (**Figure 1D**). Overall, these results suggest that MYT1L haploinsufficiency leads to a general dysregulation of apoptosis transcripts *in vitro*. Since apoptosis plays a critical role in ethanol’s neurotoxicity and may contribute to the phenotypic changes seen in our mouse model, this prompted us to investigate MYT1L’s effect on apoptosis *in vivo*.

**Figure 1.**
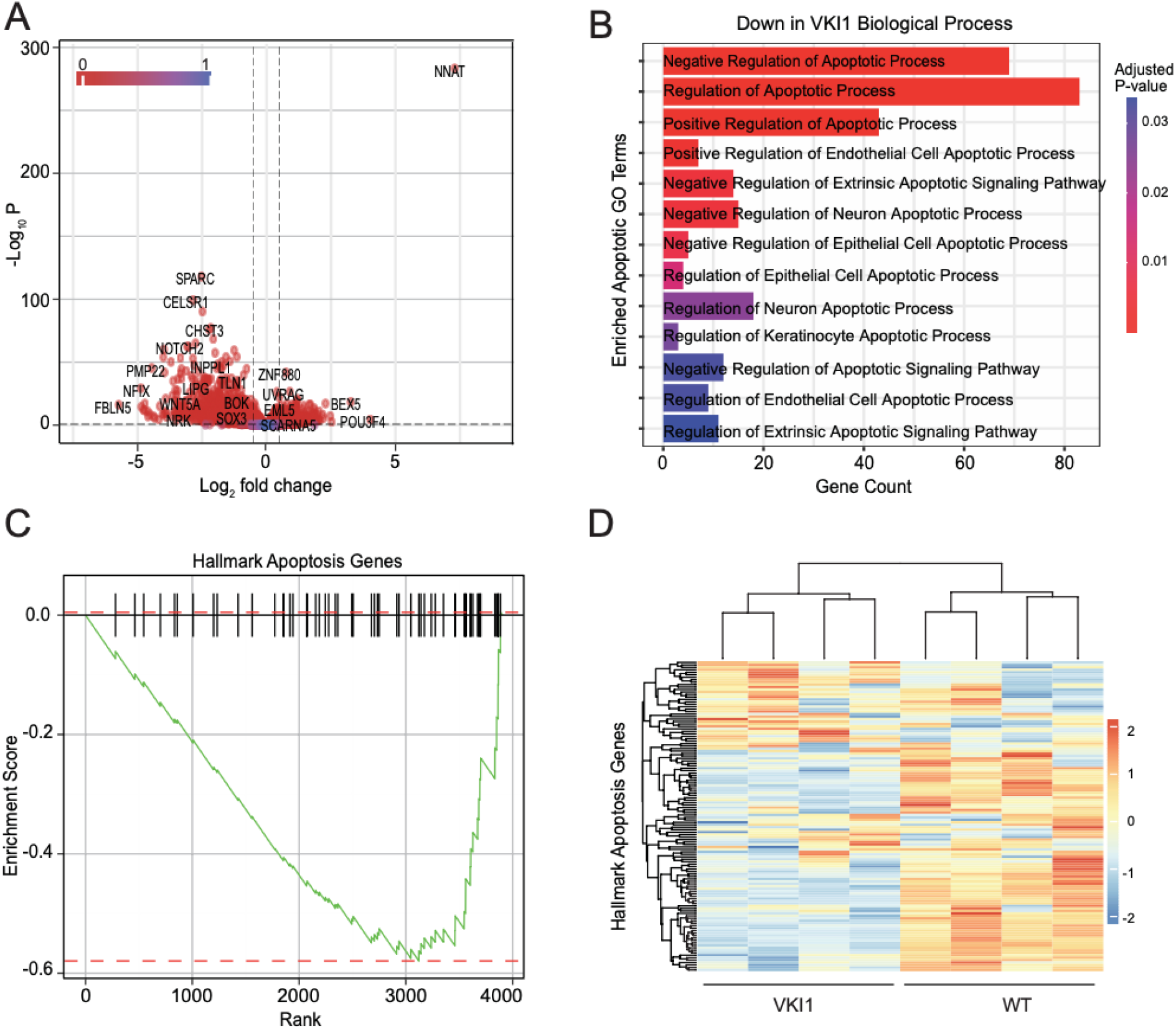
iPSC derived neurons dysregulate apoptotic programs following *MYT1L* mutation. (A) The volcano plot illustrates the differentially expressed genes identified between the wild type (WT) (control) and the *MYT1L*-S707Q variant knock-in clone 1 (VKI1) (mutant) derived from day 30 cortical interneurons. In the plot, dots represent individual genes, colored on a continuous scale by adjusted p-value. Significantly differentially expressed genes include those with an adjusted p-value < 0.05 and a fold change greater than 0.5 or less than –0.5. (**B**) Apoptosis-related biological process gene ontology (GO) term analysis for genes downregulated in the VKI1 versus WT analysis. (**C**) Gene Set Enrichment Analysis (GSEA) was conducted to evaluate apoptosis gene expression in VKI1. NES = −1.58 and adjusted p-value = 0.0026. (**D**) The heatmap shows the expression levels of hallmark apoptotic genes as in (**C**), with red = higher and blue = lower expression.

### MYT1L haploinsufficiency reduces apoptosis on P7 at moderate ethanol doses

We next examined MYT1L haploinsufficiency’s effects on apoptosis, both in unexposed Hets (compared to WT littermates), as well as in sibling ethanol-exposed mice at P7, which is the age of peak vulnerability. Het or WT mice received either ethanol (2.5 g/kg) or saline with an identical booster injection at two hours followed by perfusion 6 hours after initial injection. Brains were subsequently sectioned and immunolabeled with activated caspase-3 (AC3; a marker of cells irreversibly committed to apoptotic death) for stereological quantification. A 2-way ANOVA (Genotype x Treatment) on cerebral cell counts revealed ethanol dramatically increased apoptotic density consistent with previous research^33^ but there were no significant effects on genotype or an interaction (Treatment: F [1, 24] = 120.2, p < 0.0001, Genotype: F[1, 24] = 1.180, p = 0.2882, Interaction: F[1, 24] = 0.7729, p = 0.3880). Planned comparisons with Sidak correction revealed ethanol significantly increases apoptosis in both WT and Hets suggesting ethanol has a similar effect regardless of genotype (**Figure 2**). While stereology provides accurate estimates of apoptotic density, its use of sampling prevents calculating regional apoptotic density. We therefore quantified apoptosis on imaged sections using Object Detection machine learning (ML) software from the open source Tensorflow library (version 2.7.0) as described previously^34^. To confirm accuracy, a Pearson’s r correlation was performed between ML and stereology cerebral counts revealing a highly significant linear correlation (r = 0.9885, p < 0.0001). A simple linear regression resulted in a strong fit (R^2^ = 0.9771) and produced a best fit line slope of 1.097 indicating the counts were also similar in magnitude (**Supplementary Figure 1**). We then performed 2-way ANOVAs on ML apoptotic regional densities in the cortex, hippocampal formation, midbrain, striatum, and thalamus which all produced a main effect on treatment but not genotype with no interaction similar to stereological counts. Posthoc analyses revealed ethanol increased apoptosis in all regions for WT mice but only cortex, hippocampus, and midbrain in Hets (**Figures 2, 3A-B, Table 1**). The inability of ethanol to regionally increase apoptosis in all Het regions may suggest haploinsufficiency is partially protective against apoptosis but ethanol’s high neurotoxicity is masking this effect.

**Figure 2.**
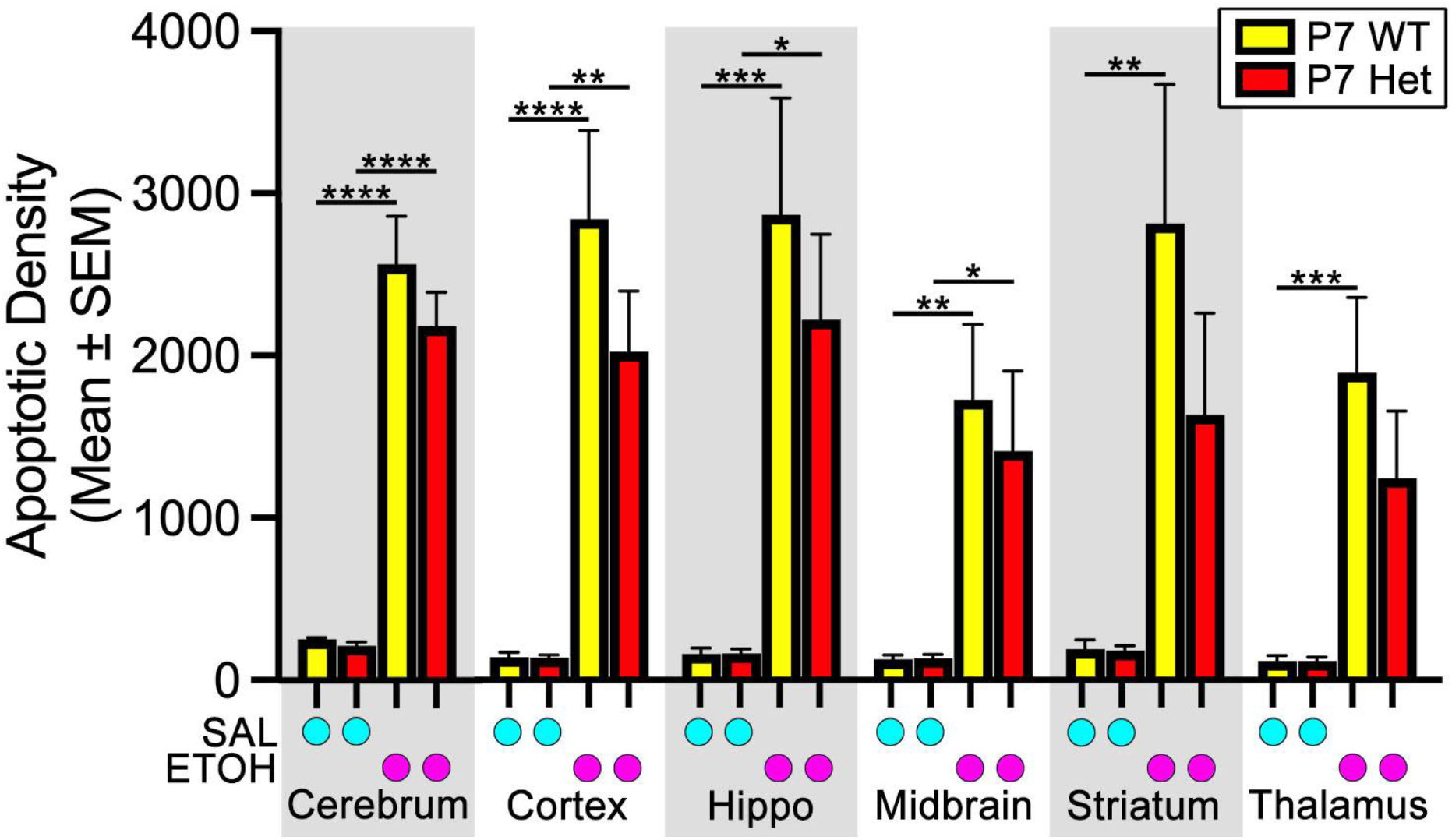
A single binge exposure to ethanol dramatically increases apoptosis on P7 but is unaffected by genotype. Mean apoptotic density in WT and Het mice following the administration of two 2.5 g/kg ethanol injections or saline (2 hrs apart) on P7. Apoptosis was quantified by stereology in the cerebrum while machine learning was performed in the cortex, hippocampus (hippo), midbrain, and thalamus. Results were analyzed with 2-Way ANOVA (see **Table 1** for main effects and interactions) followed by a planned comparisons with Sidak correction. ETOH, ethanol; SAL, saline. * p < 0.05, ** p < 0.01, *** p < 0.001, **** p < 0.0001.

**Figure 3.**
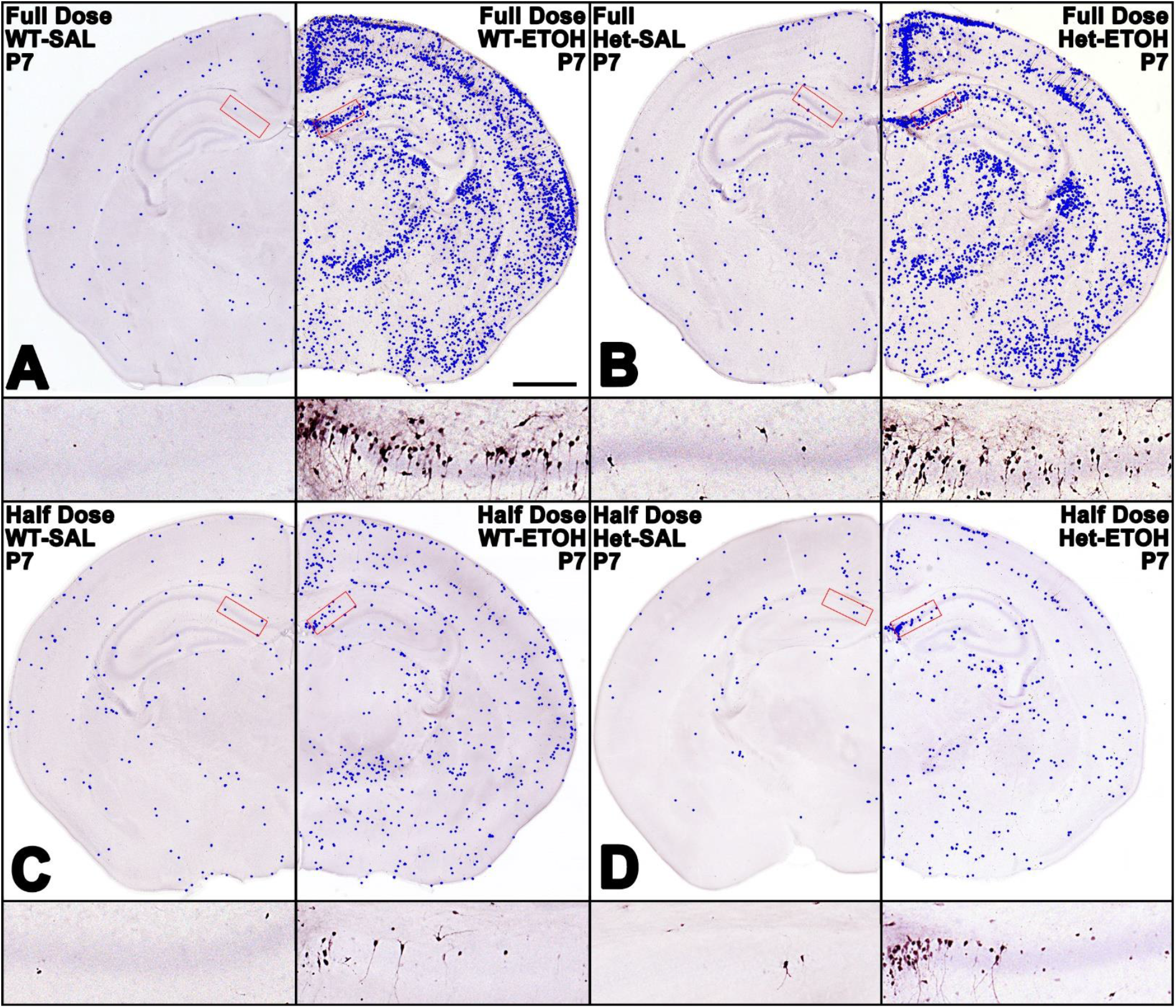
Plots of identified apoptotic cells overlaid on sections following either a full or half dose of ethanol on P7. **[A-B]** In (**A**) WT or (**B**) Het mice, representative plots of identified apoptosis using machine learning (blue dots) overlaid on top of images after two injections of saline (left) or 2.5 g/kg ethanol (right). Magnified views of outlined hippocampal CA1 regions (in red) appear below each hemisection. **[C-D]** In (**C**) WT or (**D**) Het mice, representative plots of identified apoptosis using machine learning (blue dots) overlaid on top of images after a less toxic single injection of saline (left) or 2.5 g/kg ethanol (right). Magnified views of outlined hippocampal CA1 regions (in red) appear below each hemisection. ETOH, Ethanol; SAL, saline. Scale bar = 1 mm.

**Table 1.**

| Table 1. |  |  | 2-way ANOVA (Treatment x Genotype) |  |  |
| --- | --- | --- | --- | --- | --- |
| Dose | Age | Region | Treatment | Genotype | Interact |
| Full Dose | P7 | Cortex | F [1, 24] = 41.37, p < 0.0001 | F [1, 24] = 1.320, p = 0.2620 | F [1, 24] = 1.297, |
| Full Dose | P7 | Hippocampus | F [1, 24] = 24.96, p < 0.0001 | F [1, 24] = 0.460, p = 0.5041 | F [1, 24] = 0.461, |
| Full Dose | P7 | Midbrain | F [1, 24] = 17.50, p = 0.0003 | F [1, 24] = 0.202, p = 0.6569 | F [1, 24] = 0.219, |
| Full Dose | P7 | Striatum | F [1, 24] = 12.89, p = 0.0015 | F [1, 24] = 1.098, p = 0.3051 | F [1, 24] = 1.068, |
| Full Dose | P7 | Thalamus | F [1, 24] = 20.24, p < 0.0001 | F [1, 24] = 1.013, p = 0.3241 | F [1, 24] = 1.014, |
| Half Dose | P7 | Cortex | F [1, 21] = 24.74, p < 0.0001 | F [1, 21] = 2.537, p = 0.1261 | F [1, 21] = 5.027, |
| Half Dose | P7 | Hippocampus | F [1, 21] = 48.21, p < 0.0001 | F [1, 21] = 3.824, p = 0.0640 | F [1, 21] = 9.593, |
| Half Dose | P7 | Midbrain | F [1, 21] = 22.38, p < 0.0001 | F [1, 21] = 1.371, p = 0.2547 | F [1, 21] = 3.690, |
| Half Dose | P7 | Striatum | F [1, 21] = 6.318, p = 0.0202 | F [1, 21] = 0.926, p = 0.3468 | F [1, 21] = 6.222, |
| Half Dose | P7 | Thalamus | F [1, 21] = 12.59, p = 0.0019 | F [1, 21] = 1.695, p = 0.2071 | F [1, 21] = 2.245, |
| Full Dose | P21 | Cortex | F [1, 27] = 115.1, p = 0.0001 | F [1, 27] = 6.659, p = 0.0156 | F [1, 27] = 1.852, |
| Full Dose | P21 | Hippocampus | F [1, 27] = 150.4, p = 0.0001 | F [1, 27] = 6.189, p = 0.0193 | F [1, 27] = 0.203, |
| Full Dose | P21 | Midbrain | F [1, 27] = 162.4, p = 0.0001 | F [1, 27] = 1.186, p = 0.2858 | F [1, 27] = 0.180, |
| Full Dose | P21 | Striatum | F [1, 27] = 114.7, p = 0.0001 | F [1, 27] = 8.440, p = 0.0072 | F [1, 27] = 3.518, |
| Full Dose | P21 | Thalamus | F [1, 27] = 168.1, p = 0.0001 | F [1, 27] = 4.206, p = 0.0501 | F [1, 27] = 0.056, |
| Full Dose | P21 | Dentate Gyrus | F [1, 27] = 71.11, p = 0.0001 | F [1, 27] = 2.281, p = 0.1426 | F [1, 27] = 0.280, |
| Full Dose | P60 | Cortex | F [1, 22] = 19.17, p = 0.0002 | F [1, 22] = 2.969, p = 0.0989 | F [1, 22] = 1.162, |
| Full Dose | P60 | Hippocampus | F [1, 22] = 26.37, p = 0.0001 | F [1, 22] = 1.863, p = 0.1861 | F [1, 22] = 0.482, |
| Full Dose | P60 | Midbrain | F [1, 22] = 14.72, p = 0.0009 | F [1, 22] = 2.070, p = 0.1643 | F [1, 22] = 2.434, |
| Full Dose | P60 | Striatum | F [1, 22] = 19.03, p = 0.0002 | F [1, 22] = 2.430, p = 0.1333 | F [1, 22] = 0.917, |
| Full Dose | P60 | Thalamus | F [1, 22] = 13.78, p = 0.0012 | F [1, 22] = 1.713, p = 0.2041 | F [1, 22] = 0.860, |
| Full Dose | P60 | Dentate Gyrus | F [1, 22] = 24.41, p = 0.0001 | F [1, 22] = 1.723, p = 0.2028 | F [1, 22] = 0.695, |

Grey shading indicates statistical significance

To reduce ethanol’s toxicity, we next repeated the experiment using a single 2.5 g/kg injection with perfusion 6 hours later and stereological quantification. A 2-way ANOVA (Genotype x Treatment) revealed a half-dose of ethanol produced a significant main effect of treatment (F [1, 21] = 49.41, p < 0.0001), genotype (F [1, 21] = 5.212, p = 0.0330), and a genotype-treatment interaction (F [1, 21] = 5.080, p = 0.0350). A Tukey’s posthoc analysis revealed Hets are less vulnerable to ethanol-induced apoptosis at this age possibly due to the precocious neuronal maturation described previously^27–29^. Subsequent regional quantification using ML revealed haploinsufficiency reduced density of ethanol-induced apoptosis in all regions but only reached significance in the cortex and hippocampus (**Figures 3C-D, 4, Table 1**).

**Figure 4.**
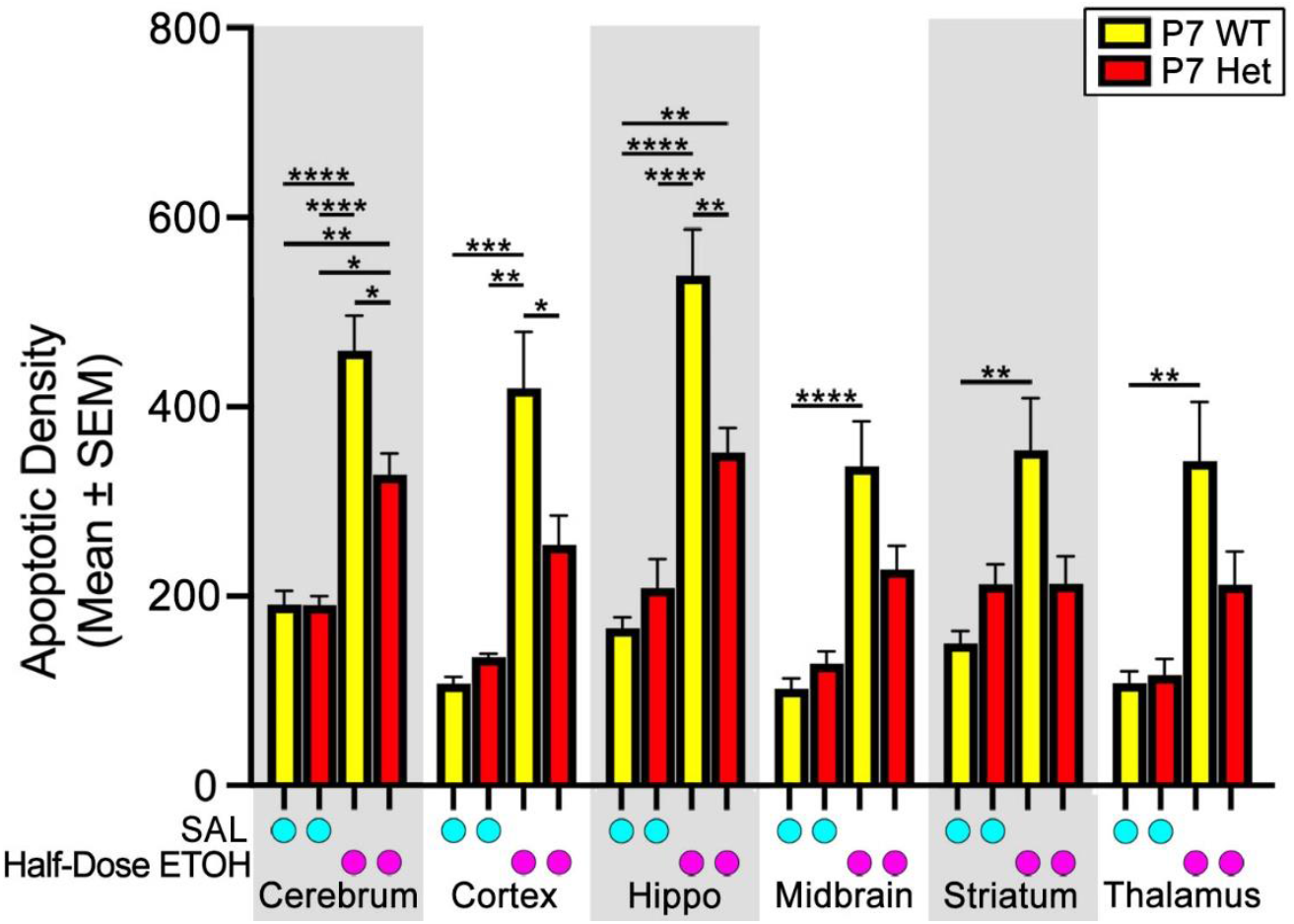
A single half-dose binge exposure to ethanol produces more moderate apoptosis on P7 that is lower in Het mice. Mean apoptotic density in WT and Het mice following the administration of a single 2.5 g/kg ethanol injections or saline on P7. Apoptosis was quantified by stereology in the cerebrum while machine learning was performed in the cortex, hippocampus (hippo), midbrain, and thalamus. ETOH, ethanol; SAL, saline. Results were analyzed with 2-Way ANOVA (see **Table 1** for main effects and interactions) followed by a Tukey posthoc comparison. * p < 0.05, ** p < 0.01, *** p < 0.001, **** p < 0.0001.

### MYT1L haploinsufficiency increases apoptosis at P21 but wild-type mice exhibit continued apoptosis in microglia and newborn dentate gyrus neurons

Vulnerability to ethanol-induced apoptosis occurs during the period of synaptogenesis (which ends around P14) and should be absent by P21^4,5^. Consistent with this, we could find no papers reporting a single binge exposure to ethanol in mice increased apoptosis in the brain at ages over P13^*35*^. We therefore examined whether haploinsufficiency extends the window of vulnerability to P21 by administering a full dose (2.5 g/kg or saline with a booster 2 hrs later) of ethanol or saline to Hets and WT littermates followed by sacrifice at 6 hours after initial injection. Quantification of cerebral apoptotic density revealed a significant main effect of treatment (F [1, 27] = 133.5, p < 0.0001), genotype (F [1, 27] = 7.309, p = 0.0117), but no genotype-treatment interaction (F [1, 27] = 2.334, p = 0.1382). While Hets (at a half dose) exhibited *reduced* apoptosis at P7, a Tukey’s posthoc revealed haploinsufficiency *increased* apoptosis at P21 (**Figures 5-6**). Surprisingly, P21 ethanol exposure increased apoptosis in WT mice more than three-fold but the apoptotic density was over an order of magnitude lower than at P7 (P7 density 2572.23 ± 296.68; P21 density 194.414 ± 19.075). In addition, the apoptotic cells’ morphology differed so much, retraining was needed for regional ML quantification using P21 tissue. A Pearson’s r correlation between stereological and retrained ML cerebral counts revealed a highly significant linear correlation (r = 0.9694, p < 0.0001), a strong fit (R^2^ = 0.9398), and produced a slope of 1.000 indicating each method produced counts very close in magnitude (**Supplementary Figure 2**). We then performed 2-way ANOVAs on apoptotic densities in the cortex, hippocampal formation, midbrain, striatum, and thalamus (**Figures 5-6)**. This revealed a main effect of treatment for all regions and a main effect of genotype for the cortex, hippocampal formation, and striatum with no significant interactions. While ethanol increased apoptosis 2-3 fold within each genotype, this effect was significantly higher in Hets within the cortex and striatum. The pattern of apoptosis also differed from P7 with apoptotic cells distributed more randomly rather than focused within a particular region or layer (**Figures 6**). In addition, in the P21 animals, the morphology of most AC3 labeling marked more punctate cells with wispy processes resembling microglia (**Figure 7B,D, Supplementary Figure 3**). Subsequent co-labeling of AC3 and microglial marker TMEM119 confirmed this cell type as microglia (**Figure 7D-F**). Interestingly, we also noticed apoptosis was occurring in the inner dentate gyrus (DG) granule layer where newborn neurons are located (**Figures 6, 7C**). DG AC3 morphology in ethanol treated animals displayed cell body and dendritic labeling which was typically absent in controls (**Figures 6, 7C,Supplementary Figure 5**). This suggested recent apoptotic death since neurodegenerative stains typically label soma and dendrites first^36^. Subsequent immunolabeling with immature granule neuron marker doublecortin and AC3 confirmed apoptosis was occurring in the inner granule cell layer where newborn granule neurons reside (**Fig 7G-I)**. No co-labeling between AC3 and proliferative marker PCNA was seen suggesting these are not proliferating neural progenitor cells (**Supplementary Figure 4**). Since this apoptosis occurred in neurons (rather than microglia), ML quantification was additionally done in the DG which revealed it contains the highest density of apoptosis of any region at this age (**Figure 5, Table 1**).

**Figure 5.**
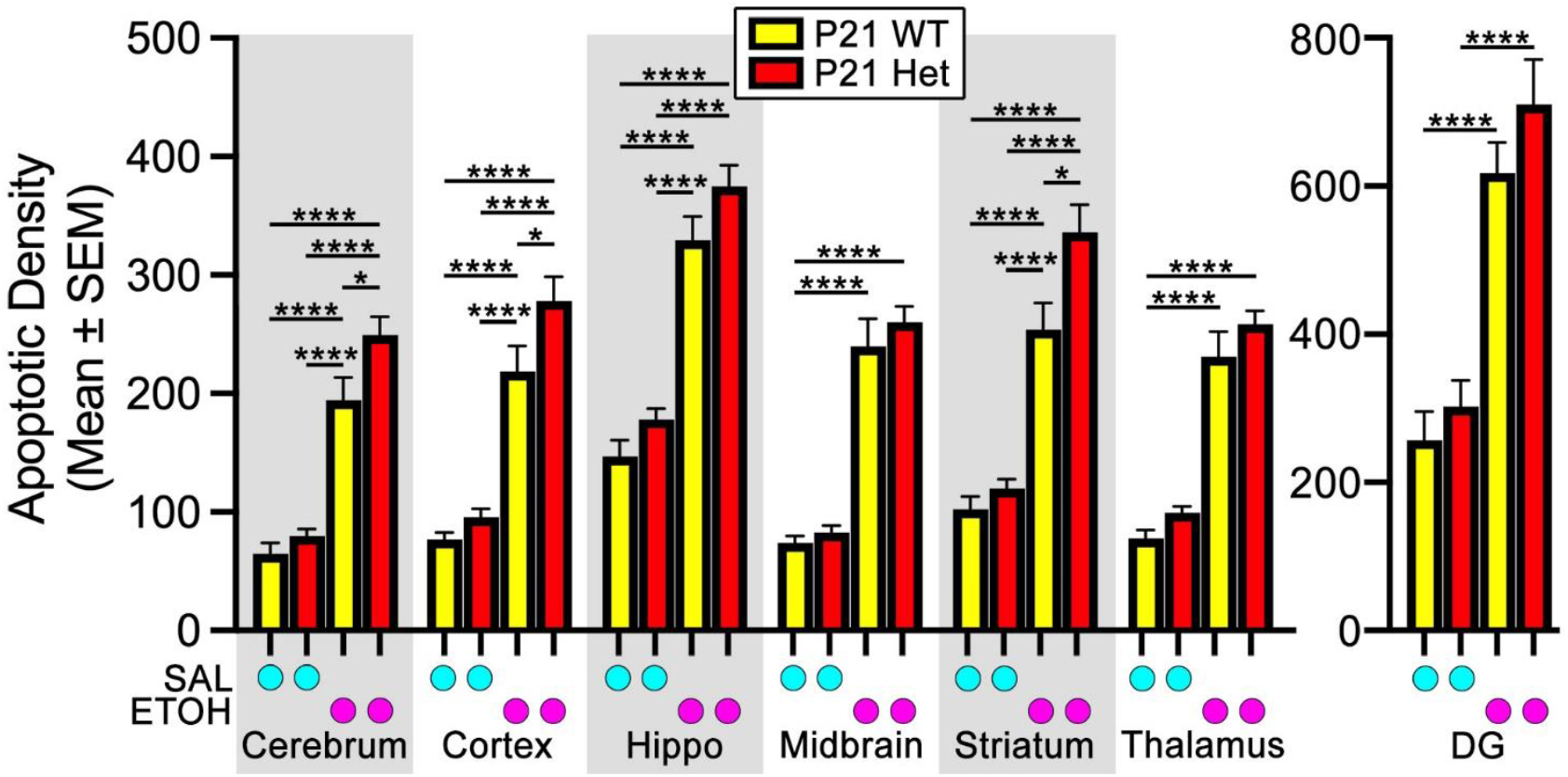
A single binge exposure to ethanol on P21 increases apoptosis in both genotypes that is increased in Hets. Mean apoptotic density in WT and Het mice following the administration of two 2.5 g/kg ethanol injections or saline (2 hrs apart) on P21. Apoptosis was quantified by stereology in the cerebrum while machine learning was performed in the cortex, hippocampus (hippo), midbrain, thalamus, and dentate gyrus (DG). The DG was quantified since it is the only area where neurodegeneration occurs at this age revealing the highest regional apoptotic density. ETOH, ethanol; SAL, saline. Results were analyzed with 2-Way ANOVA (see **Table 1** for main effects and interactions) followed by a Tukey posthoc analysis. * p < 0.05, ** p < 0.01, *** p < 0.001, **** p < 0.0001.

**Figure 6.**
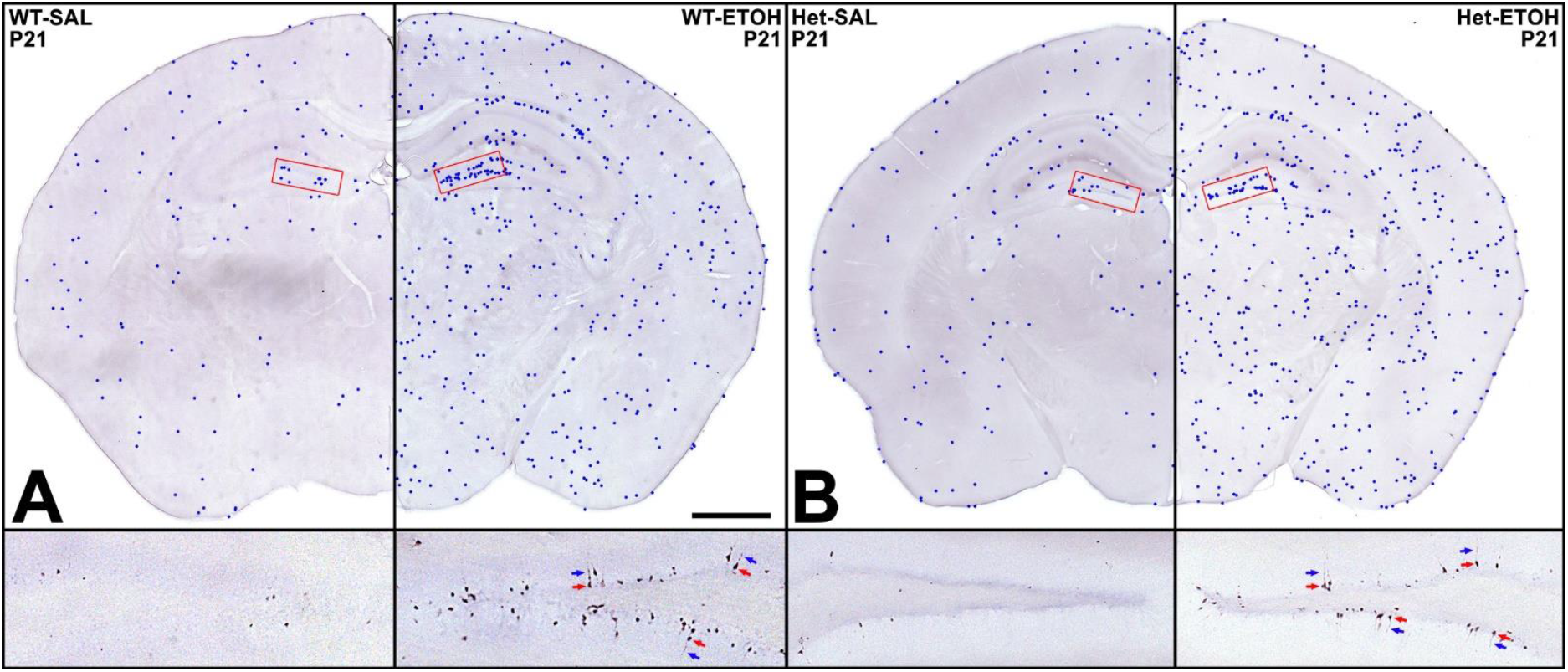
Plots of identified apoptotic cells overlaid on sections following a full dose ethanol on P21. **[A-B]** In (**A**) WT or (**B**) Het mice, representative plots of identified apoptosis using machine learning (blue dots) overlaid on top of images after two injections of saline (left) or 2.5 g/kg ethanol (right). Magnified views of outlined neurons in the dentate gyrus (in red) appear below each hemisection. Arrows point to apoptotic cell bodies (red) and dendrites (blue). ETOH, ethanol; SAL, saline. Scale bar = 1 mm.

**Figure 7.**
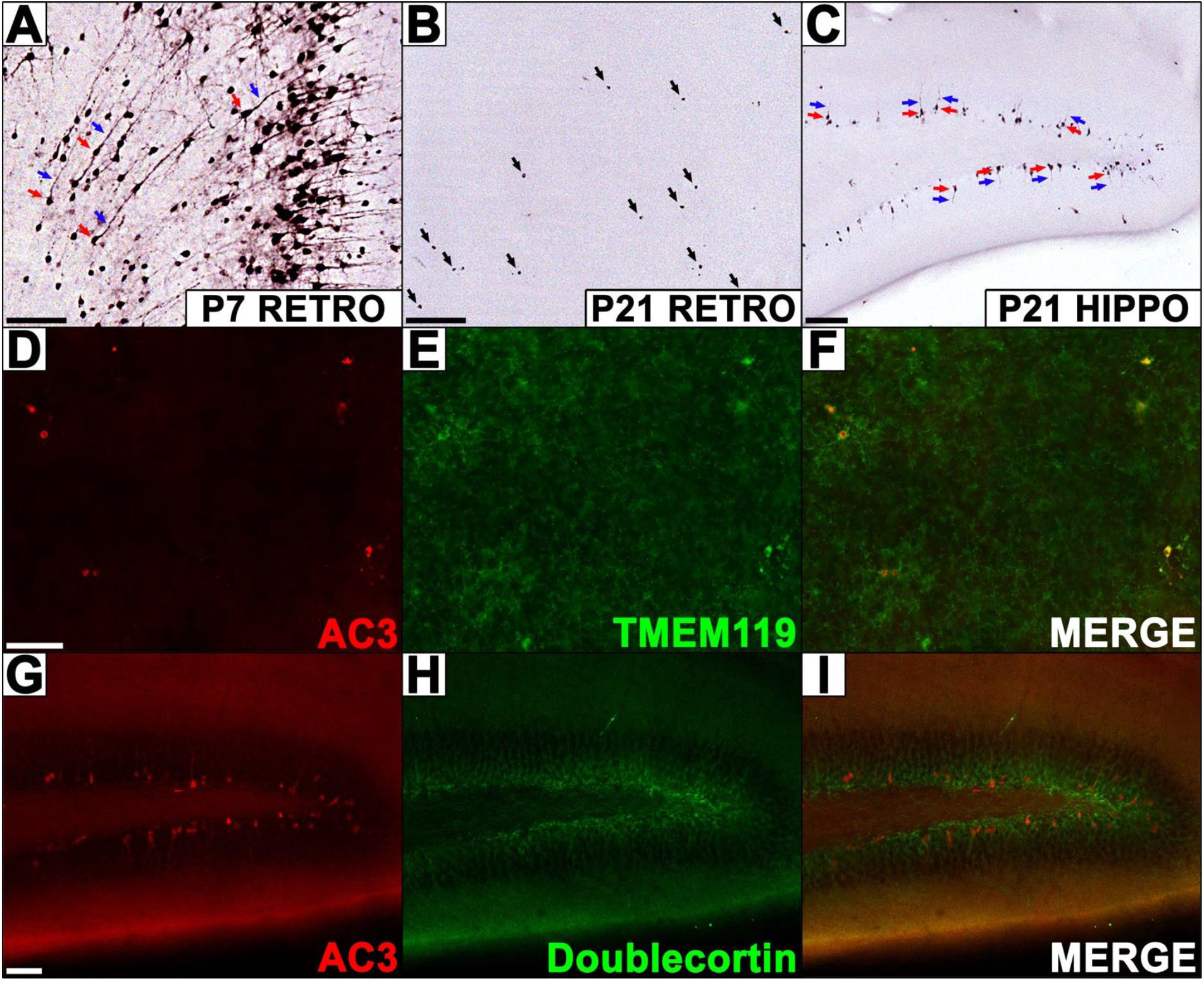
Ethanol induced apoptosis on P21 transitions to microglia and newborn granule neurons in the dentate gyrus. **[A-C]** (**A**) AC3 immunolabeling on P7 reveals heavy neuronal apoptosis in cell bodies (red arrows) and dendrites (blue arrows) in the retrosplenial cortex (RETRO). Apoptosis on P21 is either (**B**) more punctate and evenly distributed throughout the cerebrum (RETRO region pictured) or (**C**) occurs in the inner granule cell layer of the dentate gyrus where neuronal bodies (red arrows) and dendrites (blue arrows) can be seen. **[D-F]** Immunofluorescence for (**D**) AC3 and (**E**) microglial marker TMEM119 co-label (**F**) indicating microglial apoptosis. **[G-H]** Immunofluorescence for (**G**) AC3 and (**H**) immature granule neuron marker doublecortin show apoptosis occurring in this layer (**I**)confirming these are newborn granule neurons. Scale Bars: A-C = 100 uM, D-I = 50 uM.

### Adult ethanol exposure increases apoptosis in microglia and newborn granule neurons regardless of genotype

Due to continued apoptosis at P21, we next examined the apoptotic effects of ethanol in adult mice at P60. Animals were given a full dose of saline or ethanol and sacrificed 6 hours after exposure. While a significant main effect of treatment was observed (F [1, 22] = 23.19, p < 0.0001), no main effect of genotype (F [1, 22] = 2.442, p = 0.1324) or interaction was found (F [1, 22] = 2.341, p = 0.1402), suggesting that ethanol can increase apoptosis similarly within each genotype. Planned comparisons with Sidak correction to examine the effect of ethanol treatment within each genotype revealed ethanol significantly increases apoptosis in both WT and Hets (**Figures 8-9**). The morphology and regional distribution mirrored that seen at P21 indicating apoptosis continues in adulthood. Interestingly, the DG apoptotic density was comparable to other regions at this age but less than half seen at P21 for both genotypes. This is consistent with previous research suggesting adolescents are more susceptible to ethanol’s neurotoxicity than adults, which may be due to the significant reductions in DG neurogenesis as animals reach adulthood^37,38^.

**Figure 8.**
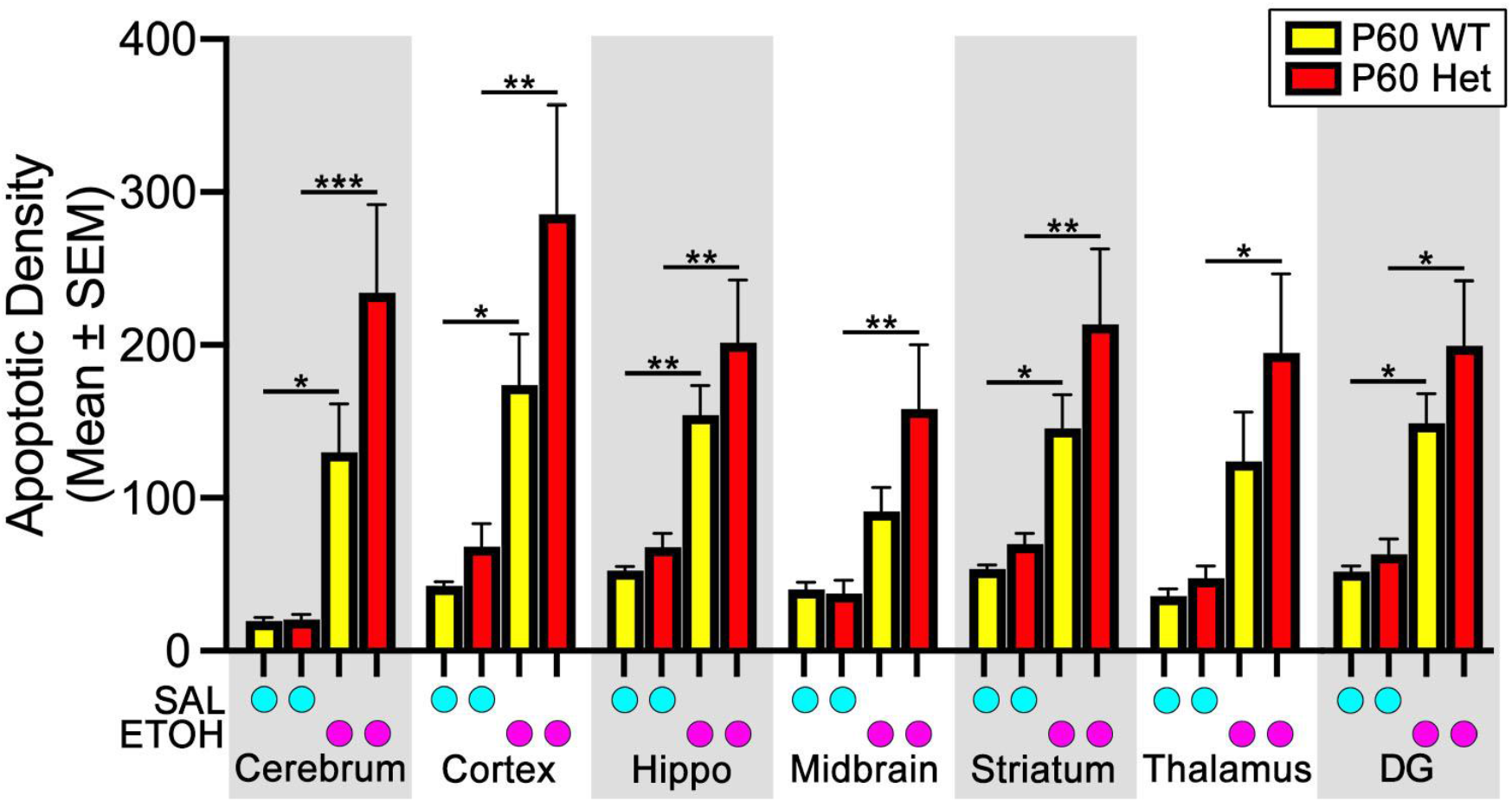
A single binge exposure to ethanol at P60 increases apoptosis in both genotypes. Mean apoptotic density in WT and Het mice following the administration of two 2.5 g/kg ethanol injections or saline (2 hrs apart) on P60. Apoptosis was quantified by stereology in the cerebrum while machine learning was performed in the cortex, hippocampus (hippo), midbrain, thalamus, and dentate gyrus (DG). ETOH, ethanol; SAL, saline. Results were analyzed with 2-Way ANOVA (see **Table 1** for main effects and interactions) followed by a Tukey posthoc analysis. * p < 0.05, ** p < 0.01, *** p < 0.001, **** p < 0.0001.

**Figure 9.**
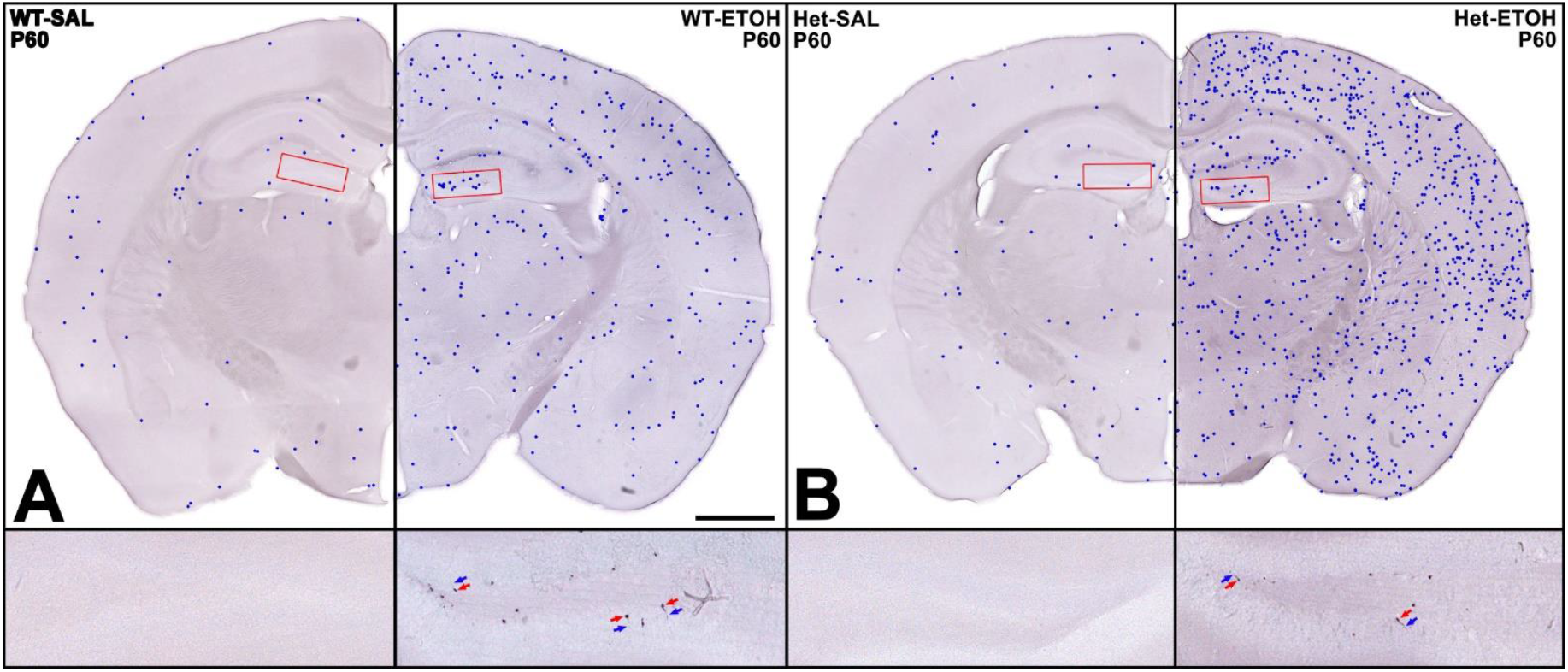
Plots of identified apoptotic cells overlaid on sections following a binge ethanol exposure at P60. Magnification in dentate gyrus. **[A-B]** In (**A**) WT or (**B**) Het mice, representative plots of identified apoptosis using machine learning (blue dots) overlaid on top of images after two injections of saline (left) or 2.5 g/kg ethanol (right). Magnified views of outlined dentate gyrus neurons (in red) appear below each hemisection. Arrows point to apoptotic cell bodies (red) and dendrites (blue). ETOH, ethanol; SAL, saline. Scale bar = 1 mm.

## Discussion

### Ethanol is neurotoxic at any age but influenced by MYT1L haploinsufficiency

Here we examined the effects of MYT1L haploinsufficiency on ethanol-induced apoptosis in the developing mouse brain. During peak vulnerability (P7), we found high ethanol exposure dramatically increased cerebral apoptosis in both genotypes. While regional counts revealed ethanol had no effect of genotype, Het apoptosis trended lower in all regions and failed to reach significance in the striatum and thalamus. This suggests haploinsufficiency may be regionally neuroprotective, but ethanol’s high neurotoxicity was overwhelming any effects. We then reduced ethanol’s toxicity by reducing the dose by half, which revealed MYT1L haploinsufficiency reduces apoptosis at this age. This may suggest the precocious neuronal differentiation (from neural progenitors) we reported previously^39^ is accelerating maturation of some neurons past their vulnerability period.

Since Het adult neurons exhibit signs of immaturity, we next examined the effects of haploinsufficiency at P21—an age where the brain is no longer supposed to be vulnerable to ethanol-induced apoptosis^4,5^. It should be noted that definitions for adolescence differ across studies, but we use the term “early adolescence” (consistent with other studies^40^) since they are mature enough to be separated from their mother^41,42^. We found ethanol increases cerebral apoptosis at P21 regardless of genotype, but the increase was significantly greater in Hets. P21 degeneration was also restricted to newly formed granule neurons in the dentate gyrus and microglia throughout the brain. Interestingly, there was no main effect of genotype in the DG (containing granule neurons), suggesting mutation only increased apoptosis in microglia.

Finally, we found ethanol increases granule cell and microglial apoptosis on P60 indicating neurotoxicity continues into adulthood, but genotype had no influence. Compared to apoptotic density at P21, microglial counts were slightly reduced but there was a dramatic decrease in the DG (less than half) which is likely due to the significant reductions of neurogenesis seen as mice reach adulthood^37^.

### Myt1l *mutation has moderate effects on ethanol’s neurotoxicity*

In the absence of ethanol, it did not appear that the *Myt1l* mutation strongly influenced neuronal apoptosis in the developing brain, suggesting physiological apoptosis is not affected during neurodevelopment. Importantly, even in the presence of a factor that increases neuronal apoptosis during brain development (ethanol at P7), *Myt1l* mutation actually decreased the amount of neuronal death. One possible explanation is that the *Myt1l* mutant neurons have matured precociously and have more rapidly moved beyond the window of vulnerability to apoptosis. Indeed, transcriptional studies suggest a dysregulated maturation trajectory across age in *Myt1l* mutants^27,30,43^.

Interestingly, the same mutants showed increased apoptotic response to ethanol at P21, but in microglia not granule neurons. Notably, microglia do not express MYT1L in the mice used in this study^27,43^. Thus, the increased apoptosis in these cells must be a non-cell autonomous effect. A study in a different mouse model of *Myt1l* mutation suggested microglial transcriptional activation, especially in adults^44^. It may be that some features of unusual neuronal function in *Myt1l* mutants may induce microglia to be in a proliferative or otherwise activated state that may make them more vulnerable to ethanol-induced apoptosis. However, this same microglial signature was not apparent in the adult transcriptomes of the mouse line used here, either in cortex ^27^, or hypothalamus^45,46^. Nonetheless, future studies targeting microglial states across the various mutants in parallel may be warranted.

### Ethanol produces apoptosis in newly formed dentate gyrus granule neurons

Translated to human use, our regimen of a single acute binge exposure would be far more common than chronic multiday ethanol exposures used in most preclinical research and may have important implications for the estimated 61.4 million people (21.7% of the population) age 12 and older who binge drink in the United States^47^. It may also provide insight into alcohol-related brain damage produced by chronic use since there is controversy about what toxicity is produced directly by ethanol verses secondary factors such as nutritional deficiency, excessive stimulation during withdrawal, and compensatory changes to the brain^10,48–50^. One benefit of our acute experimental design is that it isolates ethanol’s direct neurotoxicity which occurs before secondary factors have time to emerge. For instance, ethanol’s ability to reduce hippocampal neurogenesis is well-known and intensely studied due to the ever-expanding roles neurogenesis is thought to play in learning and memory and neurologic/psychiatric diseases such as depression, Alzheimer’s disease, and epilepsy ^50,51^. While others have found DG degeneration following more chronic ethanol exposures lasting days to weeks^11,52–55^, there is debate as to whether it is caused by ethanol’s direct effects or secondary factors produced by chronic use^10,48– 50,56^. Interestingly, the conclusions from these studies is usually that cell death is necrotic^11,57,58^ but there are compensatory changes producing neurodegeneration such as excitotoxicity (a type of necrotic death due to overstimulation of a neuron^59^) during withdrawal or thiamine deficiency^56,60^. Importantly, AC3 detects apoptosis before these secondary effects can emerge and does not detect necrotic degeneration^61^. Additionally, our proposed mechanism (see below) is through prolonged inhibition rather than excitation. Another outstanding question is whether reduced neurogenesis is caused by changes in proliferation, differentiation, migration, or cell survival^56,62^. Our findings show granule cell apoptosis is at least one major cause of this phenomenon.

We suggest this apoptosis occurs through the same mechanisms that regulate neuronal apoptosis during the synaptogenesis period on P7. Adult newborn granule neurons are susceptible to apoptosis during two phases. The first occurs within four days of cell birth and kills the majority of immature granule neurons (neuroblasts)^63^. The second occurs about three weeks after neuronal birth when the cell sends axonal projections to integrate into local circuitry (i.e., the neuron’s synaptogenesis period)^38,64^. During this critical window, conditional knockout of NMDA receptor function produces apoptosis which implies survival is use dependent (i.e., stimulation from synaptic connectivity prevents apoptosis)^63,64^. It also suggests the NMDA antagonism produced by ethanol may simulate poor synaptic connectivity and induce granule neuron apoptosis similar to what occurs during the P7 synaptogenesis period. If true, ethanol may have a magnified effect by deleting the few granule neurons that have established synaptic connectivity and would have otherwise been integrated into hippocampal circuitry.

### Ethanol is acutely neurotoxic in microglia

While numerous studies have examined the effects of ethanol on microglia, they generally focus on microglial activation producing neurotoxic reactive oxygen species and neuroinflammatory factors^8,56,65^. As a result, our finding that ethanol induces rapid apoptosis in microglia reveals a surprising and previously unknown neurotoxic effect of ethanol. Interestingly, since chronic ethanol use reduces brain volume that recovers with abstinence, scientists have suspected ethanol produces microglial toxicity due to their adult regenerative capacity (something that applies to DG granule neurons also)^50,66^. Consistent with this, diffusion tensor imaging has revealed increased grey matter mean diffusivity (MD) in people with alcohol use disorder which was replicated in rodents following a chronic month-long ethanol exposure^67^. While increased MD is typically attributed to neuroinflammation, rodent histology revealed a highly significant reduction in microglia in every region but no effect on astrocytes. Administration of a selective microglial toxin also increased MD suggesting reductions in this cell type were producing imaging changes. Another imaging study examined alcohol-dependent subjects during withdrawal and found markers for microglial activation are lowered. While the authors predicted withdrawal to produce microglial activation, they suggested this effect was caused by reduced microglial density^68^. These studies are consistent with the microglial reductions reported in postmortem brains from individuals with alcohol use disorder^66,69^. Taken as a whole, both preclinical and clinical research suggest the acute microglial neurotoxicity reported here can lead to prolonged reductions with chronic alcohol use. Microglia are best known as the resident macrophages in the brain that regulate immune response, phagocytosis, and inflammation. However, they are also responsible for synaptic pruning which, if interrupted, can alter synaptic transmission, functional connectivity, and inhibit forgetting^65,70,71^. They also regulate adult hippocampal neurogenesis by phagocytizing apoptotic neurons and are thought to promote DG granule neuron proliferation, differentiation, and survival^72^. As a result, ethanol’s ability to kill newborn granule neurons and microglia may have a magnified effect on the hippocampal function.

### Summary

Previous preclinical research has found ethanol produces rapid apoptosis within hours at P7 but this vulnerability was thought to rapidly disappear by P14^*4*,*35*^. Motivated by our finding that *Myt1l* mutations transcriptionally alter apoptotic regulators, we examined this gene’s effects on ethanol-induced apoptosis at different ages. We discovered that this gene can moderately influence vulnerability in a complex age- and cell type-specific manner with reduced neuronal apoptosis at P7, increased microglial apoptosis at P21, but no effect at P60. It is notable that this animal model is sensitive to even moderate effects and may provide a novel robust platform to study ethanol’s neurotoxicity moving forward. In early adolescent and adult mice, we also discovered the previously unknown ability of a single binge ethanol exposure to produce apoptosis within hours in newborn hippocampal granule neurons and microglia. Translated into human use, this suggests ethanol-induced neurotoxicity is far more common than previously known and possibly affects the 1 in 5 people that binge drink in the United States^47^. These acute effects may also contribute to long-term structural and functional deficits associated with chronic alcohol use and highlight critical cellular targets that warrant further investigation.

## Methods

### Mouse Models

All mouse procedures were approved by the Institutional Care and Use Committee at Washington University School of Medicine. Mice were bred and housed in the vivarium at Washington University in St. Louis. Animal husbandry conditions consisted of translucent plastic cages with corncob bedding, *ad libitum* access to lab diet and water, a 12 hour light and dark cycle, and standard room temperatures (20-22°C) and humidity (50%). *Myt1l* Het mice have been described previously^27^. Breeding pairs of *Myt1l* Het and wild type C57BL/6J mice (JAX Stock No. 000664) were used to generate male and female *Myt1l* Het and WT littermates, with genotyping conducted as described^27^.

### Procedures

The WT and *Myt1l* Het mice were administered either ethanol or saline at postnatal days 7 (P7), P21, or P60. Ethanol (2.5 grams/kg) or saline was administered intraperitoneally at 10 μL per gram weight plus a second injection two hours later followed by perfusion six hours after the initial injection. Some P7 mice received a half dose where a single ethanol 2.5 grams/kg or saline dose was administered with perfusion at six hours. Mice were deeply anesthetized using Fatal Plus and transcardially perfused with 4% paraformaldehyde in PBS. The cerebrum of each animal was serially sectioned in the coronal plane using a vibratome set at 75 μM. A representative set of every eighth section was collected and immunolabeled for activated caspase-3 (1:1000, Cell Signaling Technology, Cat#: 9661L); allowing quantification of cells irreversibly committed to apoptotic death. For immunolabeling, free floating sections were quenched in 3% hydrogen peroxide in absolute methanol for 10 minutes, immersed for 1 hour in a blocking solution (2% Bovine Serum Albumin, 0.2% Dry Milk, 0.8% TX-100 in PBS), and incubated overnight at room temperature with the AC3 primary antibody in solution (1:1000, Cell Signaling, Cat#: 9661L, 1% Bovine Serum Albumin, 0.4% TX-100 in PBS). The next morning, sections were incubated with a biotinylated secondary antibody solution (1% Bovine Serum Albumin in PBS), reacted with an avidin-biotin conjugate kit (Vectastain Elite ABC Kit, Cat#: PK-6100, Vector Labs), and visualized using the chromogen VIP (VIP Substrate Kit; Cat#: SK-4600, Vector Labs). For immunofluorescence, detection of apoptotic immature granule neurons, microglia, and proliferating cells was achieved using doublecortin (1:3000, Santa Cruz Biotechnology, Cat#: sc-271390), TMEM119 (1:400, Synaptic Antibodies, Cat#: 400 004), and PCNA (1:2400, Cell Signaling, Cat#:2586S) respectively co-labeled with AC3. All immunofluorescent images were taken using a Leica DM6B running Leica Application Suite X (v3.8.1.26810) with a Leica DFC7000 T digital camera.

### Stereological Analysis

Apoptotic cells were stereologically counted using Stereoinvestigator Software (v 2019.1.3, MBF Bioscience, Williston, Vermont, USA) running on a Dell Precision Tower 5810 computer connected to a QImaging 2000R camera and a Labophot-2 Nikon microscope with electronically driven motorized stage. A rater, blind to treatment, traced each hemisection and stereologically quantified the number of apoptotic cells using the unbiased optical fractionator method. This information was then used to estimate the density of apoptotic cells per cerebrum. In one animal, a large amount of AC3 positive blood vessel staining made quantification difficult and was eliminated from future analysis.

### Quantification of apoptotic cells using machine learning

To obtain regional counts, machine learning was used to generate apoptotic cell plots using the open source Tensorflow software library (version 2.7.0) and a suite of custom written Python (version 3.9) scripts as described previously^34^. Quantification was performed on a Windows 10 personal computer with a RTX 3080 graphics card (64GB DDR4-3200 PC4-25600 RAM, and AMD Ryzen 9 3950X 16-Core 3.5 GHz CPU). After AC3 immunolabeling, section photocomposites were produced by imaging at 10X with a Leica DM 4000B microscope equipped with a Leica DFC310FC camera using Surveyor software (Version 9.0.2.5, Objective Imaging, Kansasville, WI). Images were saved in TIF format and manually adjusted (blind to genotype) using Photoshop CS5.1 by white point leveling to produce a white background. A python script then applied an unsharpmask filter (PIL library: Amount 75%, Radius 2.4, Threshold: 0) to each image. Since Tensorflow cannot process large files, each photocomposite was sliced into smaller jpg images (256 × 256 pixels). Slices adjacent to others had an additional 100 pixel guard zone added to that side which overlapped its neighbor. This ensured identified objects were not partially cut off by the edge of the slice. If an object was identified in the guard zone, only objects whose bounding box center was on the half of the zone closest to the slice were counted. Around 150 slices were randomly selected to create a training set for machine learning. A human manually outlined objects in each slice using LabelImg software (https://github.com/heartexlabs/labelImg) and saved annotations in XML format. Slices containing no imaged tissue or no apoptotic cells were not used and replaced with another random slice. The images and their annotations were then randomly divided into a training and testing set (containing approximately 100 and 50 slices respectively) and converted into the TFRecord format for training. Machine learning was then performed using the Object Detection script in the Tensorflow Library and run for 105,000 steps for P7 images or 150,000 steps for P21/P60 images (trained models are available upon request). The resulting trained model was then used to identify objects (both apoptotic microglia and neurons) in the rest of the slices using a threshold of 0.001 (P7 images) or 0.1 (P21 and P60 images) and results exported for further analysis. For P7 mice, a python script was used to filter out false positives by analyzing pixels converted to greyscale within the bounding box of each identified cell. Cells where the difference between the 95th and 5th percentile pixel intensity was below 150 were eliminated. This removed cells where its bounding box contained uniform pixel intensity. For P21 and P60 images, bounding boxes whose area overlapped by 80% or more had the object with a lower score removed to eliminate double counting. The remaining counts were then used to generate cell plots and regionally quantify apoptosis. All python scripts used in this study are available upon request, free of charge, from Washington University for non-commercial, non-clinical use.

### Statistical Analysis

Data was presented as the mean ± standard error of the mean (SEM). A 2-way Treatment x Genotype ANOVA was performed on apoptotic density (GraphPad Prism 10.5.0) for each region. A Tukeys posthoc was performed when a main effect of treatment and either an interaction or genotype was significant. A planned comparisons with Sidak correction was performed to examine the effect of treatment within each genotype when only a main effect of treatment was detected. For Pearson’s r comparisons of cerebral counts machine learning counts were compared to stereological hemisection counts that were doubled before comparison.

## Data Availability

RNA-seq data was examined from previously published work and can be found at the Gene Expression Omnibus (GEO) at the following accession: GSE244189. All downstream RNA-seq analysis was performed in R with RStudio. No unique code was generated.

## Acknowledgements

This work was supported by the IDDRC (P50HD103525) and R01MH124808 to KLK.

## Supplementary Figures

**Supplementary Figure 1.**
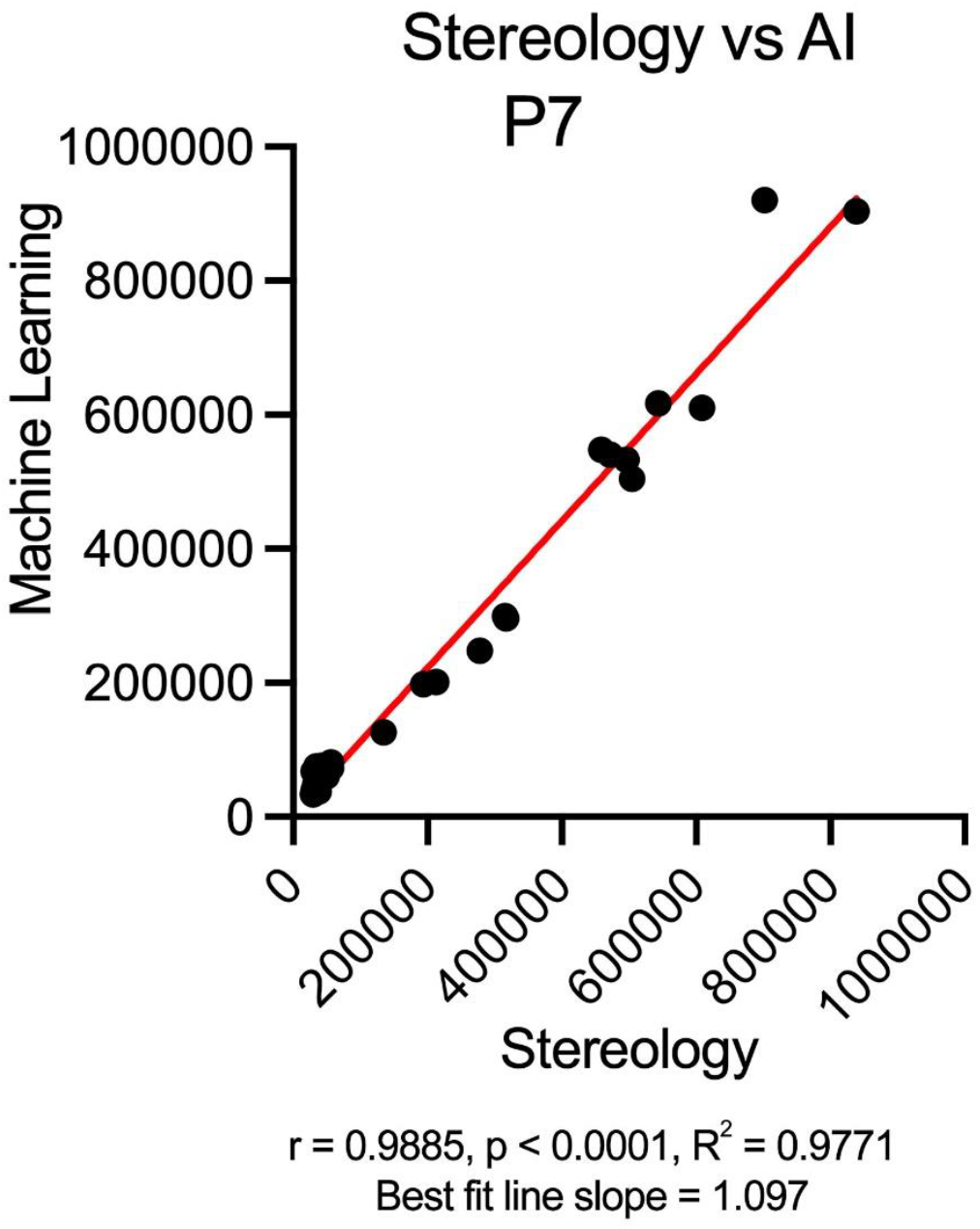
The estimated number of ethanol-induced apoptotic cells on P7 using stereology and machine learning are correlated and similar in magnitude. A Pearson’s r correlation of cerebral apoptotic cells quantified using machine learning and stereology on P7 revealed a highly significant linear correlation. A best fit line revealed a slope of 1.097 indicating correlated counts were also similar in magnitude.

**Supplementary Figure 2.**
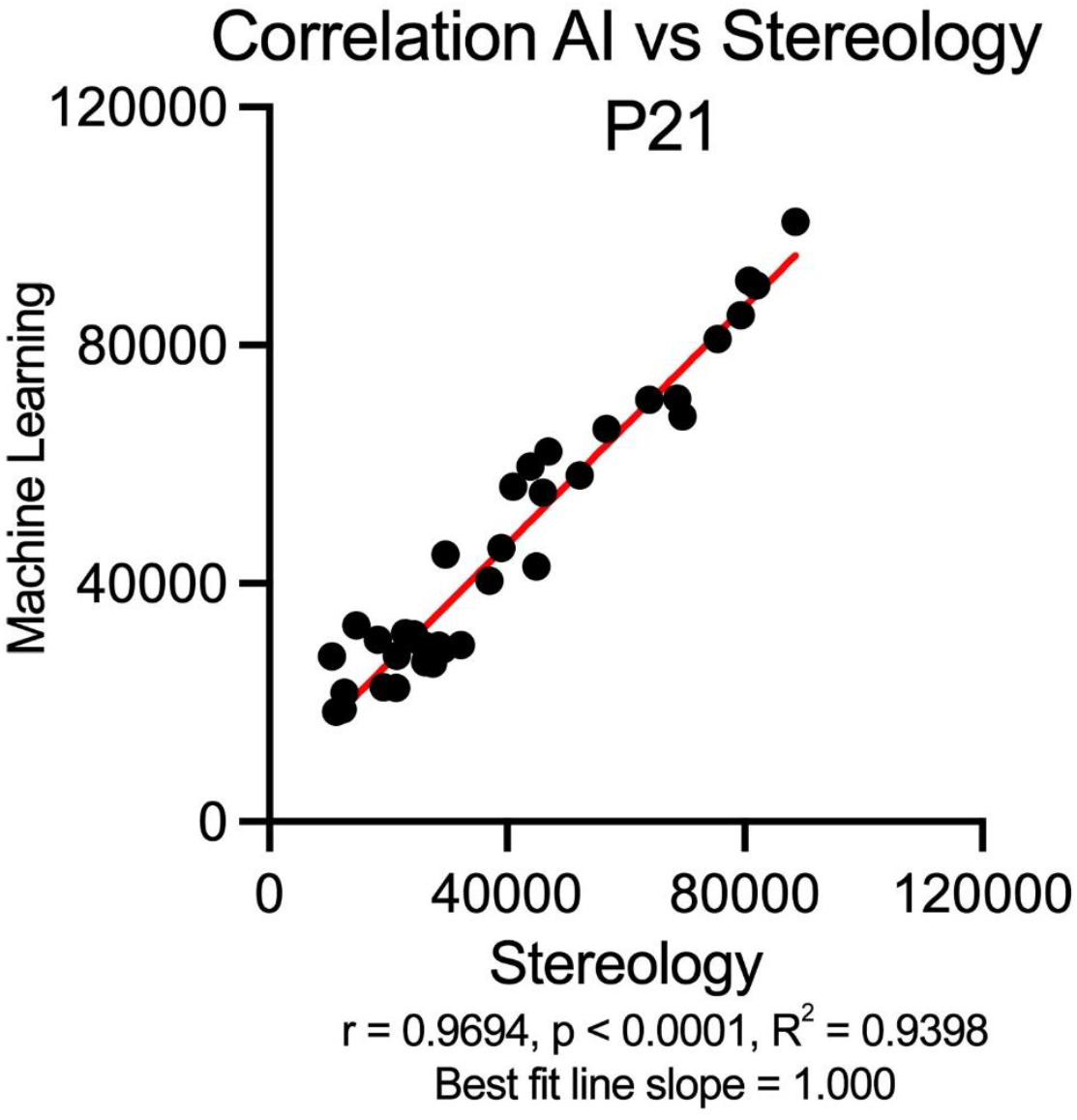
The estimated number of ethanol-induced apoptotic cells on P21 using stereology and machine learning are correlated and similar in magnitude. The P7 and P21 apoptotic morphology differed so drastically, a second machine learning training run was required. A second Pearson’s r correlation of cerebral apoptotic cells quantified using machine learning and stereology on P21 revealed a highly significant linear correlation. A best fit line revealed a slope of 1.000 indicating correlated counts were also similar in magnitude..

**Supplementary Figure 3.**
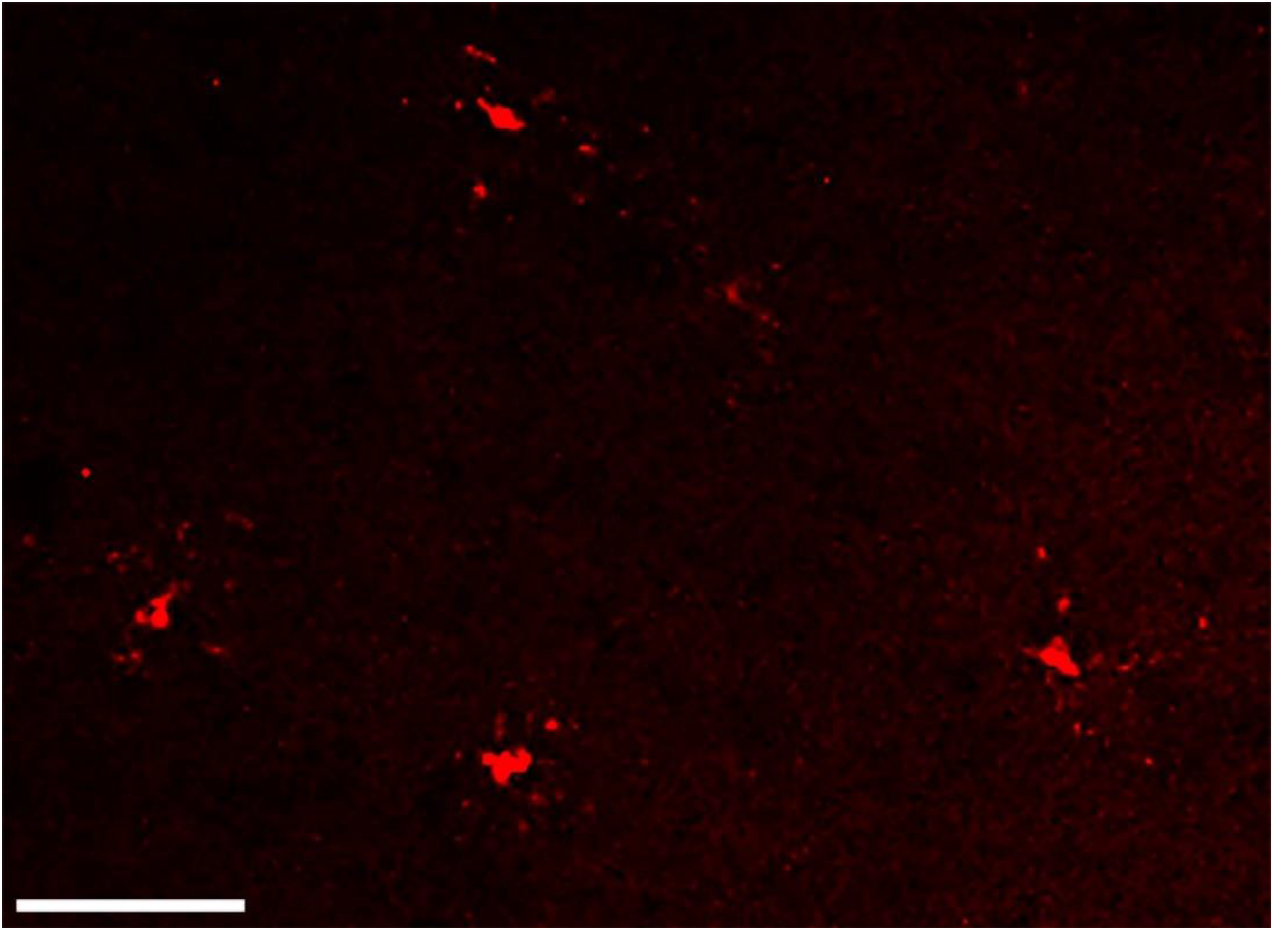
AC3 positive cells outside the hippocampus resemble microglia. Outside the dentate gyrus, AC3 labeling of apoptotic cells revealed staining of a central mass often surrounded by wispy processes similar to microglia. Scale bar = 50 uM.

**Supplementary Figure 4.**
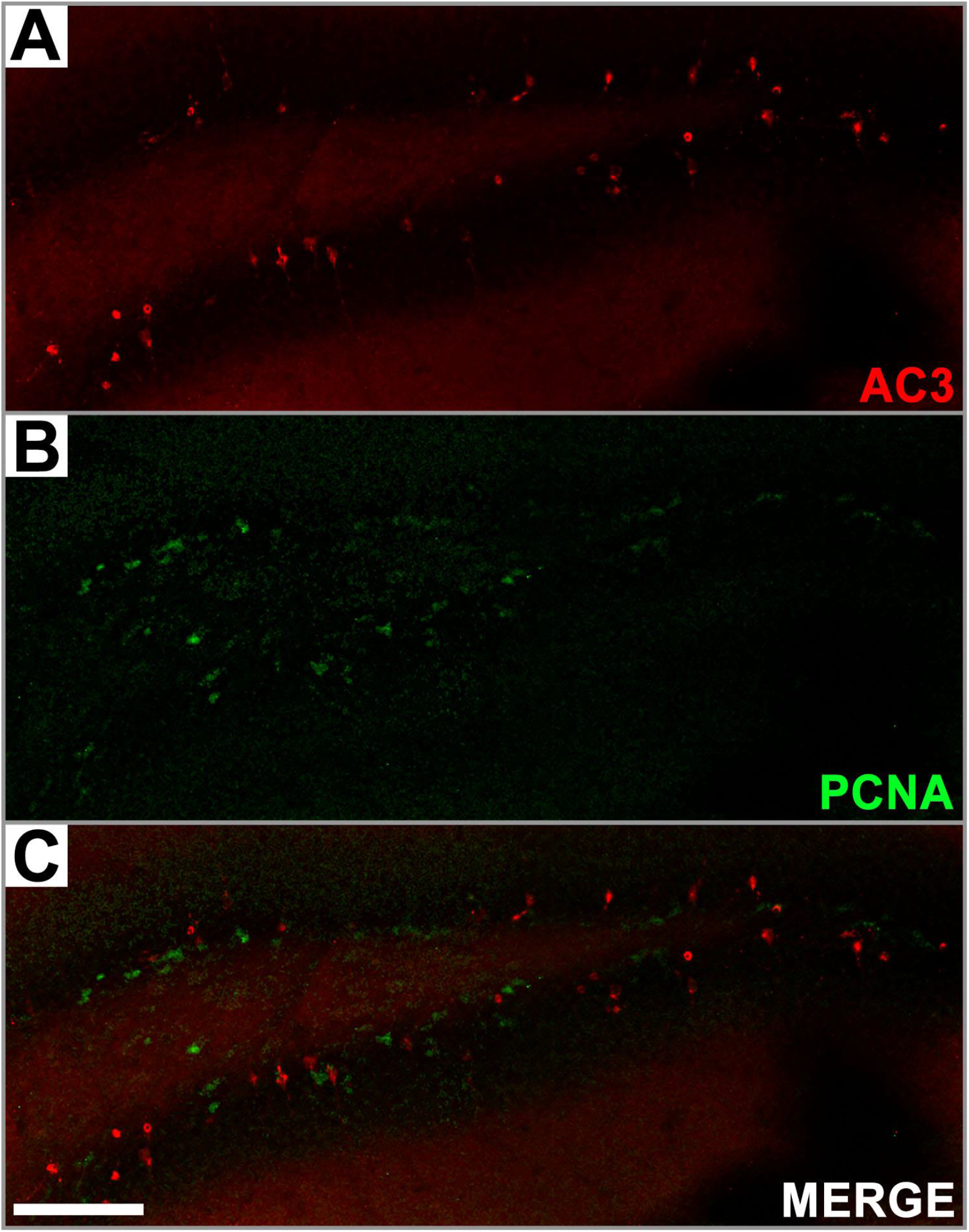
Dentate gyrus apoptotic cells are not proliferating neural progenitor cells. Immunofluorescent labeling with AC3 and the proliferative marker PCNA reveal no co-labeling indicating apoptotic cells are not proliferating neural progenitor cells. Scale bar = 100 uM.

**Supplementary Figure 5.**
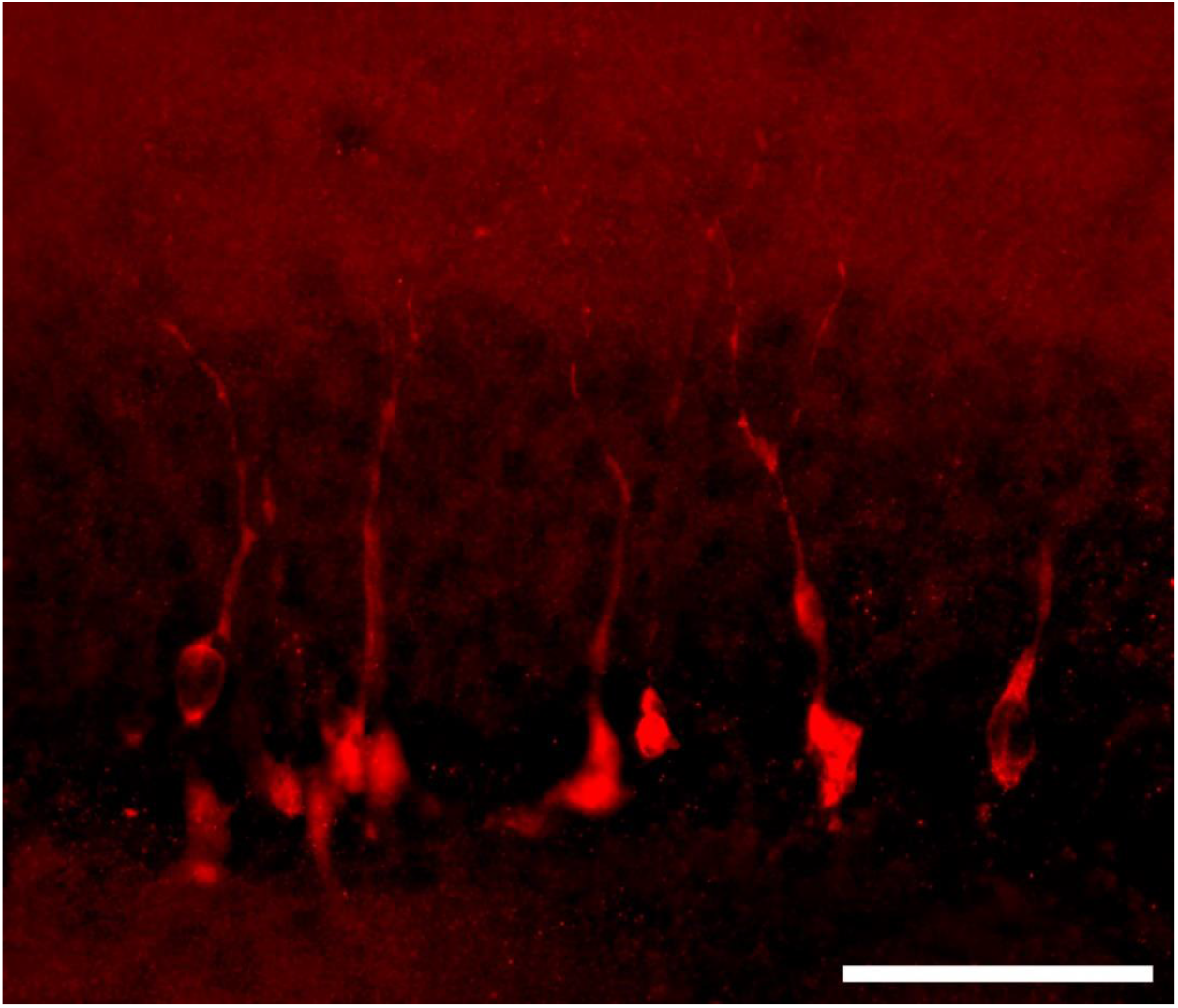
AC3 immunolabeling of apoptotic granule neurons marks soma and dendrites. Apoptotic granule neurons in the dentate gyrus exhibit AC3 labeling of soma and dendrites which extend dorsally to the perforant path. Scale Bar = 50 uM.

